# Using Oxford Nanopore Technology direct RNA sequencing to identify depurination events induced by ricin and other ribosome inactivating proteins

**DOI:** 10.1101/2021.08.13.456275

**Authors:** Yan Ryan, Abbie Harrison, Hannah Trivett, Catherine Hartley, Jonathan David, Graeme Clark, Julian A. Hiscox

## Abstract

Depurination is a frequent modification to both DNA and RNA, in DNA causing point mutations through misincorporation, in RNA, disabling ribosomes and halting protein synthesis. Some modifications of nucleic acids can be determined by direct sequencing using Oxford Nanopore Technologies (ONT). However, the identification of modifications is often limited by noise and their variety and number. Ricin is a toxin which enters cells and depurinates an adenine base in the sarcin-ricin loop of the large ribosomal subunit. This leaves only a ribose backbone, thus inhibiting protein translation. In humans, biological threat agents and ribosome inactivating proteins, such as ricin and saporin, depurinate base 4605 on the 28S rRNA providing a single defined target to try and identify. We postulated that the depurination event could be detected using ONT direct RNA sequencing through a change in charge in the ricin loop. A software tool was developed, RIPpore, that quantified the adenine modification from direct RNA sequencing data of ribosomal RNA purified from respiratory epithelial cells exposed to ricin. This provided a novel method of directly identifying ricin exposure and a basis for ONT’s utility in detecting lesions in nucleic acids caused by depurination events.

## Introduction

Deciphering the depth and breadth of the epigenome and epitranscriptome has been an ongoing challenge, with 17 DNA^1^ and 173 RNA^2^ modifications described to date in the literature. Current methods to detect these changes are modification specific, such as bisulphite sequencing for 5-methylcytosine^3^ or antibody based immunoprecipitation and sequencing of N(6)-methyladenosine^4^. Oxford Nanopore Technology (ONT) sequencing, by measuring the ionic current and dwell time of nucleotides traversing through a nanopore, can potentially capture the entire spectrum of modifications present on non-amplified DNA or RNA. Base calling software (basecallers) take a signal representing a nucleotide or sequence of nucleotides from a sequencing device and translate this into the constituent nucleotides in a format that can be analysed. ONT have developed basecallers for Nanopore sequencing such as Guppy, that translate ionic current and dwell time into the nucleotides AGCT/U. In contrast to other sequencing methodologies, the nucleic acid being sequenced on an ONT flow cell is analysed across the nanopore, five nucleotides at a time (denoted as a kmer). ONT basecallers initially used hidden Markov models to predict and designate each base, whereas later iterations utilise neural networks for increased accuracy. Newer versions of Guppy can infer 5mA and 6mC in DNA. One critical factor has been to create reference samples featuring modifications at known positions. However, this has proved technically challenging. Currently, no ONT base callers have yet been trained to recognise modifications that are not common and especially those on RNA.

Depurination is one such modification, where either an adenine or guanine base is enzymatically or chemically cleaved from the phosphodiester backbone, leaving a gap in the DNA or RNA strand. Depurination spontaneously occurs in DNA at a rate of 2,000-10,000 events per day per cell^5^, which are broadly resolved by base excision repair^6^. However mis-repairing can occur, which leads to mutation and in some cases oncogenesis^7^. Depurination of RNA, depending on context, has multiple downstream effects, from halting translation^8^, preventing packaging of viral RNA into capsids^9^, or halting protein synthesis in ribosomes by cleavage of a key catalytic adenine^10^.

Ribosome-inactivating proteins (RIPs) are a class of toxin which inhibit protein synthesis via the depurination of a specific adenine base present within a conserved ribosomal region known as the sarcin-ricin loop (SRL)^11^. RIPs induce irreversible hydrolysis of the N-glycosidic bond at adenine 4605 (A4605) in *Homo sapien* 28s rRNA^12^ (this cleaved site is often referred to as A4324, referencing the corresponding *Rattus norvegicus* homologue). A4605 depurination is an ideal candidate for identification via direct RNA sequencing, as this results in the modification of a single known target site. Ricin is one of the most well-known RIPs due to potent cytotoxic properties and is a significant risk to both human and animal health; with many deliberate exposures having been reported, including the assassination of Georgi Markov, a Bulgarian dissident writer^13, 14^. Ricin is a plant lectin and holotoxin composed of two subunits individually referred to as ricin toxin A chain (RTA) and ricin toxin B chain (RTB). The B chain binds to the cell surface and facilitates entry whilst the A chain catalyses depurination, classifying ricin as a type II RIP. Related type I RIPs, such as saporin, lack this B chain and therefore require the use of an artificial cell entry mechanism (i.e., transfection reagent) to enter a cell and cause toxicity.

To investigate whether depurination of A4605 in the SRL could be identified by direct RNA sequencing, a lung epithelial cell line (A549) was exposed to ricin. Post-exposure, direct RNAseq was used to evaluate the ricin induced signal alterations in 28s rRNA. The data indicated that depurination of A4605, formed a signal analogous to what would occur if the base were U4605. This sequence transposition was exploited as a proxy measure for depurination. RIPpore, a Python script was developed which leveraged the transposition to quantify and isolate depurinated reads, enabling signal level investigation of wholly modified and unmodified populations. The methodology presented herein provided a novel approach to identify SRL depurination in a sample exposed to RIPs. This opens a door to the future transcriptome or genome wide mapping of apurunic sites via ONT direct sequencing and the detection of the effects of a biological/chemical threat agent.

## Results

To identify depuration, a combined biological exposure, sequence and computational strategy was developed. Cells were exposed to ricin and saporin (two RIPs) and the resulting RNAs analysed by direct RNAseq using an ONT flowcell (Fig. 1).

**Fig. 1.**
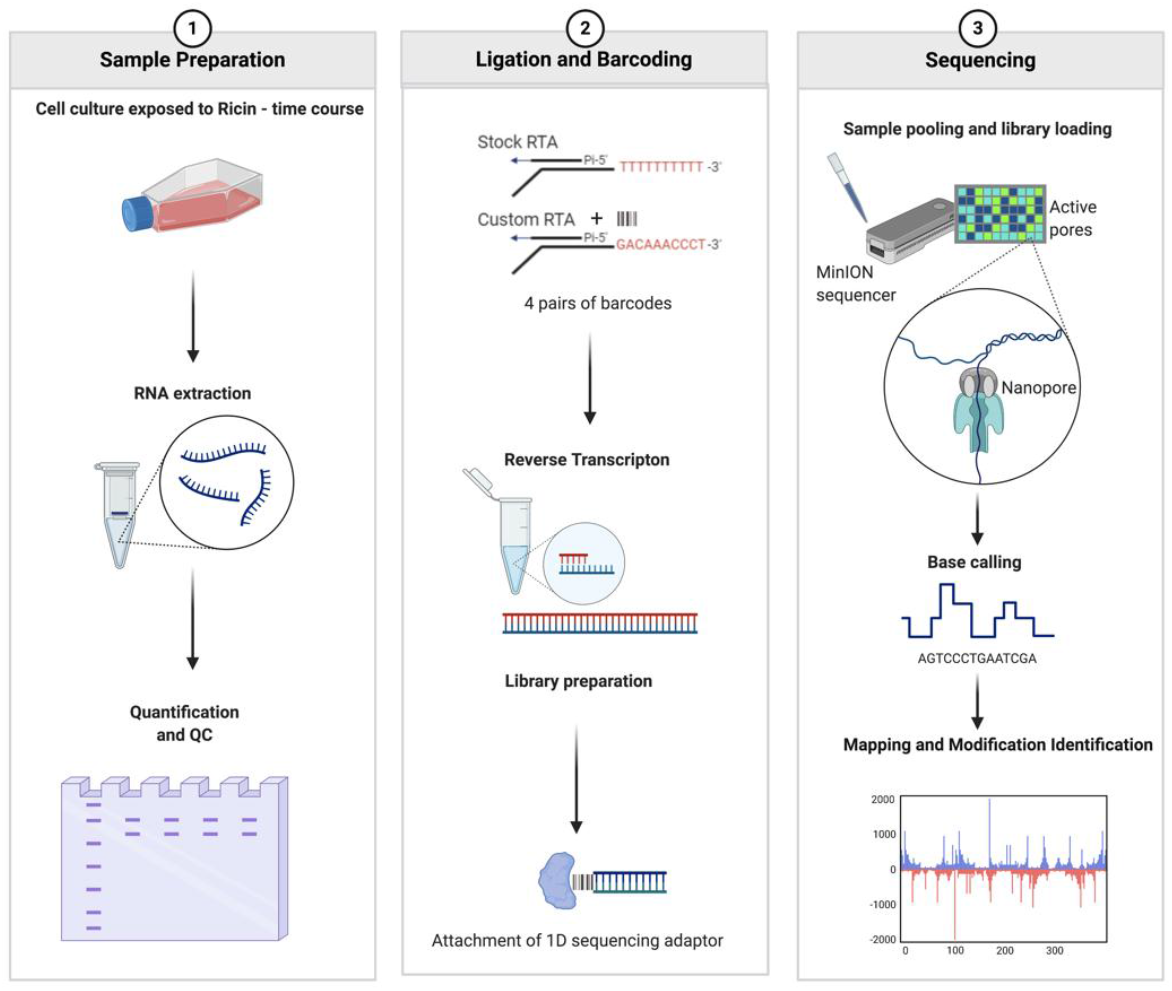
Strategy for assaying the effects of ricin using a direct RNA sequencing approach. A549 cells were exposed to ricin and total RNA extracted from cells and purified in a continuous procedure. A select portion of rRNA potentially containing the A4605 modification was directly sequenced on a flow cell. Software was developed, RIPpore, that took the output data and quantified the adenine modification.

### Ricin purification and cytotoxicity assay in cell culture

The ricin toxin used in this study, was isolated from the *Ricinus communis* (*subsp. Zanzibariensis*) plant and underwent a two-stage purification process from bean, to crude, to purified preparation. Purification of ricin was confirmed by Coomassie blue staining, with solvent extracted crude preparation showing multiple products, and a single product of the expected molecular weight after HPLC purification (Fig. 2A). To assess the toxicity of the crude and purified ricin, a WST-1 cytotoxicity assay was performed on the cell lines of choice - A549 cells (Fig. 2B). (Note the complete Commassie blue staining pattern is shown in Supplementary Fig. 1). Cell viability plateaued at concentrations of ricin above 10 pM, and further increases in concentration did not decrease cell viability, suggesting these were saturating amounts. The half maximal effective dose (EC50) had only a 2 fM difference between preparations. To ensure saturation and to provide parity to other methodologies^15^ a 1 nM concentration of purified ricin was used throughout this study. Purified ricin toxin was also used to reduce the possibility that effects were not due to other constituents of castor beans.

**Fig. 2A.**
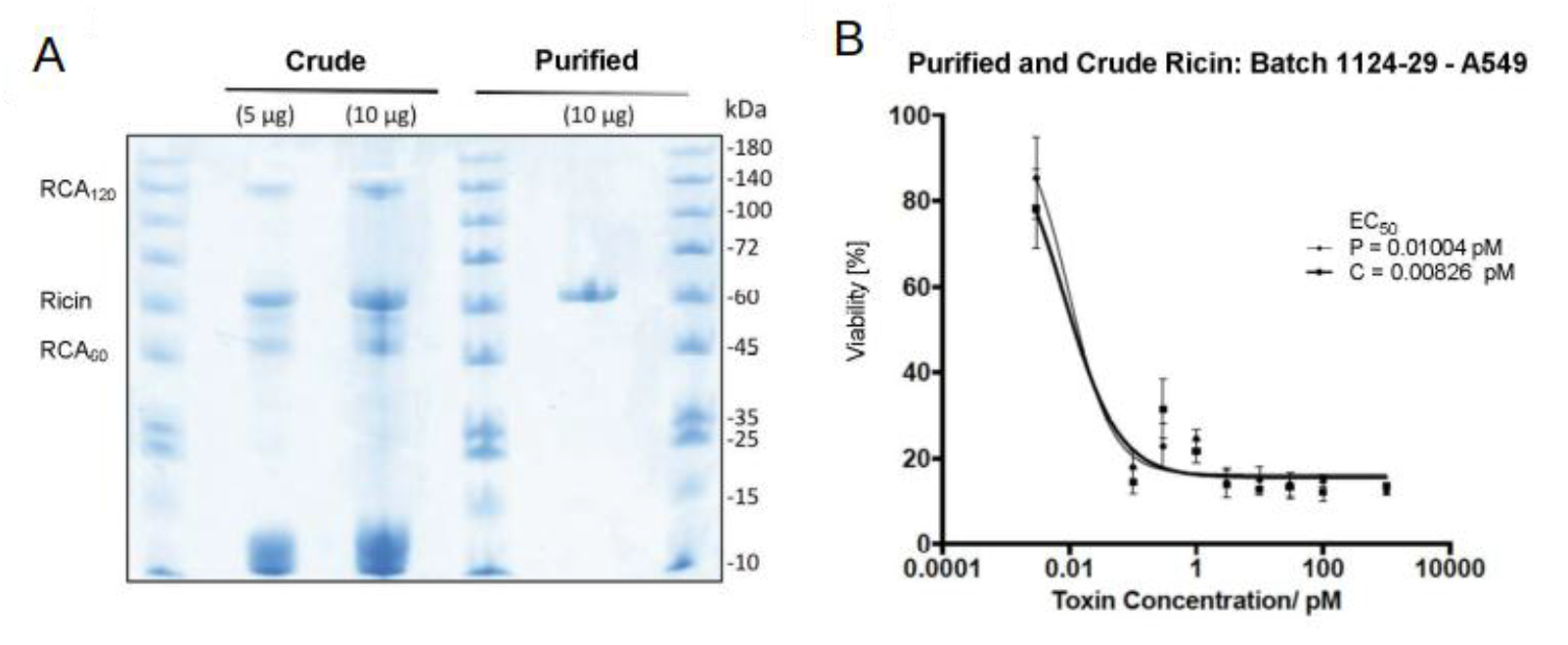
Coomassie blue analysis of crude and purified ricin toxin preparations. Two different amounts of the crude preparation were evaluated (5 μg and 10 μg) and a single preparation of the purified fraction (10 μg). Molecular weight markers are indicated to the right and the expected size of purified ricin and the R. communis agglutinins (RCA120 and RCA60) are indicated to the left. (B). WST-1 viability assay on A549 cells exposed to increasing concentrations of crude (C) and purified (P) ricin toxin for 48 hours showed a plateau in toxicity beginning at 10 pM and a difference in EC50 of 2 fM.

### 28s Ribosomal RNA targeted Oligo Design

To directly sequence the depurinated base(s) in the SRL, ONTs Reverse Transcription Adapter (RTA) Oligonucleotides A and B (designed to ligate to polyA-tailed mRNAs) were customised to specifically ligate to non-polyadenylated ribosomal RNA. The 3’ polyT run was replaced with the reverse complement of the final 10 3’ bases of the 28s rRNA (producing: GACAAACCCT). For flow cell efficiency, barcoding was used to multiplex samples. Supplementary Table 1. describes the number of reads and samples per flow cell, Supplementary Table 2. shows an overview of per sample read count, coverage of position 4605 in the 28s rRNA and percentage of reads which mapped to the 28s rRNA. Once ligated, samples were reverse transcribed, generating a cDNA hybrid for enhanced stability and to minimise secondary structures in the 28s RNA, facilitating the transit of the native RNA molecule through the nanopore for sequencing.

### RIP induced depurination caused a shift in charge intensity at the apurinic site to be identified as an uracil

Direct RNAseq using Oxford Nanopore enabled the study of chemical modifications on RNA by measuring potential alterations in charge. The hypothesis was tested that depuration of adenine 4605 by a RIP would lead to a detectable signal change or miss-base calling. To test this hypothesis, hACE2-A549 cells were exposed to ricin or saporin (a type I RIP), total RNA extracted and sequenced directly. Reverse transcription was performed to aid the sequencing of the 28S. Guppy was used to call the sequence of the 28S rRNA moving through the pore. We predicted that if the apurinic 4605 site was present, this would be potentially miss base called, whereas an unmodified base would be correctly called as an adenine.

Direct sequence reads were mapped to human 28s rRNA using minimap2 (although uracil was basecalled as thymine, this is referred to as an uracil residue). At 2 hours post-ricin exposure, 12% of nucleotides at the site of depurination (A4605) had an apparent change from adenine to uracil compared to the equivalent sequence from unexposed cells (Table 1). To quantify and further study this in the multiple samples, a python script was developed, RIPpore, to count the type and number of each nucleotide occurring at base 4605. By analysing the unexposed controls and cells exposed to ricin B chain (RTB), a baseline level of noise was established with ∼ 0.3% of reads called as uracil (Table 1). This identified a limit of detection of the approach, whereby RIP induced uracil transpositions above a cut-off of 0.4% were reported as depurinated. This cut-off was calculated by taking the maximum percentage plus standard deviation of uracil bases called at position A4605 in control and RTB exposed cells. Updating Guppy from version 3.0.6 to 4.4.2, and repeating base calling on control data led to a decrease in the noise of uracil from ∼1.5% (data not shown) to a mean of 0.277 (Table **Error! Reference source not found**.). Rebasecalling data from RIP exposed cells, found no change in uracil calls (data not shown). An increase in apparent depurination events was also observed in 28s rRNA purified from cells exposed to saporin.

**Table 1:** Table of descriptive statistics of percentage of reads called as uracil by Guppy

| Exposure | Number<br>of<br>Samples | Min | Max | Median | Mean | Standard<br>Deviation |
| --- | --- | --- | --- | --- | --- | --- |
| Control | 3 | 0.21 | 0.336 | 0.284 | 0.277 | 0.063 |
| B-Chain | 4 | 0.15 | 0.171 | 0.159 | 0.159 | 0.0012 |
| Saporin | 3 | 7.546 | 16.97 | 8.524 | 11.015 | 5.184 |
| Ricin 2<br>hours | 3 | 9.177 | 15.34 | 11.65 | 12.036 | 3.129 |
| Ricin 4<br>hours | 3 | 24.765 | 29.71 | 26.48 | 25.983 | 3.995 |
| Ricin 6<br>hours | 3 | 26.54 | 35.06 | 30.66 | 30.749 | 4.262 |
| Ricin 24<br>hours | 3 | 19.78 | 39.64 | 23.81 | 27.743 | 10.5 |

Boxplots were used to visualise counts of each nucleotide at base 4605 across all samples. The data showed a time-dependent increase in depurination, by concomitant increase in uracil and decrease in adenine (with significance determined by pairwise t-tests) (Fig. 3). The proportion of cytosine and guanine both increased in count to around 3% each at this site. The depurination effect of ricin was detected at the initial time point analysed (2 hours post exposure), peaked at 6 hours, and by 24 hours had declined. All ricin exposures showed a significant difference at this position from the negative control or RTB.

**Fig. 3.**
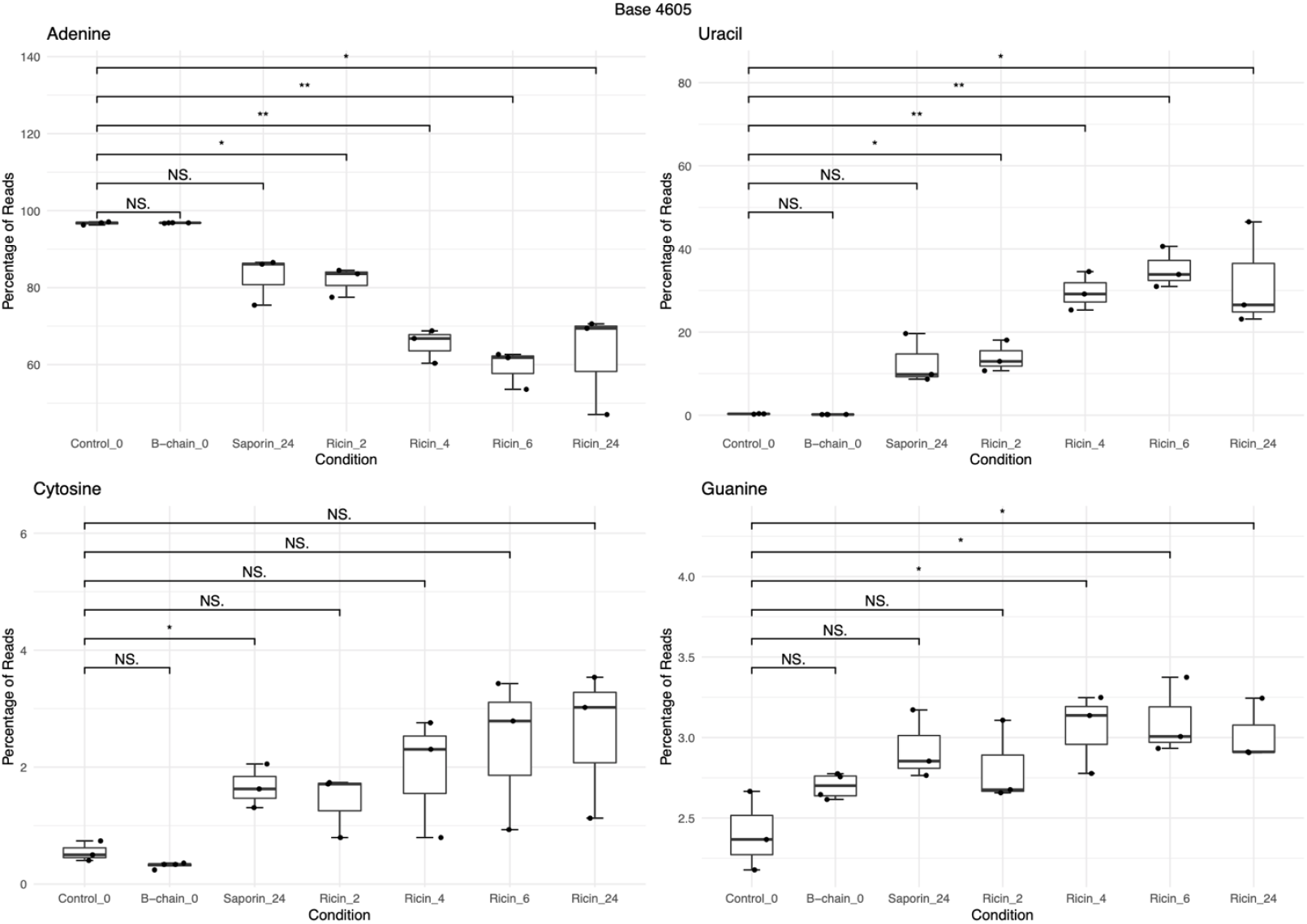
Boxplots showing percentage of nucleotide composition at position A4605 in 28S rRNA from A549 cells either a control, b-chain or saporin at 24 hr or ricin at 2, 4, 6 and 24 hours post exposure. Pairwise t-tests assessing control to each exposure were used to perform significance testing. The decrease in base called adenine was inverse to the increase in the uracil calls. Cytosine and guanine showed an average of 3% increase from control.

### Charge intensity analysis of the ricin loop shows a shift caused by depurination

As variation in raw measurements such as dwell time and charge intensity are utilised by base calling algorithms to predict nucleotides, the influence that depurination in the 28s rRNA had upon these raw features was measured. Previous studies suggested sequencing of depurinated DNA resulted in an increase in dwell time using solid state (not ONT) nanopores^16^. The charge intensity and dwell time at a per base resolution and the changes induced by depurination were compared between the different exposures and controls and were visualised using Nanocompore. The data indicated that there was a decrease in charge associated with the apurunic site, with the proportion of reads with this shift increasing with time, in RNA from cells exposed to either ricin or saporin when compared to the unexposed controls (Fig. 4). Nanocompore identified a distinct increase in charge density formed below 107 mean intensity units (Nanocompores unit), which was absent in the control reads and increases in a time dependent manner. This effect was seen initially as a small peak in the exposed samples, which increased in size over time to a distinct peak, with a concomitant decrease in the canonical peak at 115 mean intensity units. Ricin exposure for 2, 4, 6 and 24 hours (Figs. 4A, B, C and D) respectively, caused a rise in reads at a lower charge density, increasing from 2-6 hours, peaking, and plateauing between 6 hours and 24. The 2 hour exposure (Fig. 4B), had the lowest significance in difference to control of the ricin exposures (Figs. 4G and H), with each subsequent exposure increasing in significance. Exposure with ricin B chain only, had little discernible difference in charge density at position 4605, but a slight increase in dwell time (Figs. 4E and G). Exposure of cells to saporin (Fig. 4F), resulted in an increase in reads with a lower charge density and a slight increase in dwell time, which was greater than those observed in cells exposed to B-chain only. Ricin exposure produced a noticeable and significant increase in lower charge reads at site 4605, in a time dependent manner.

**Fig. 4.**
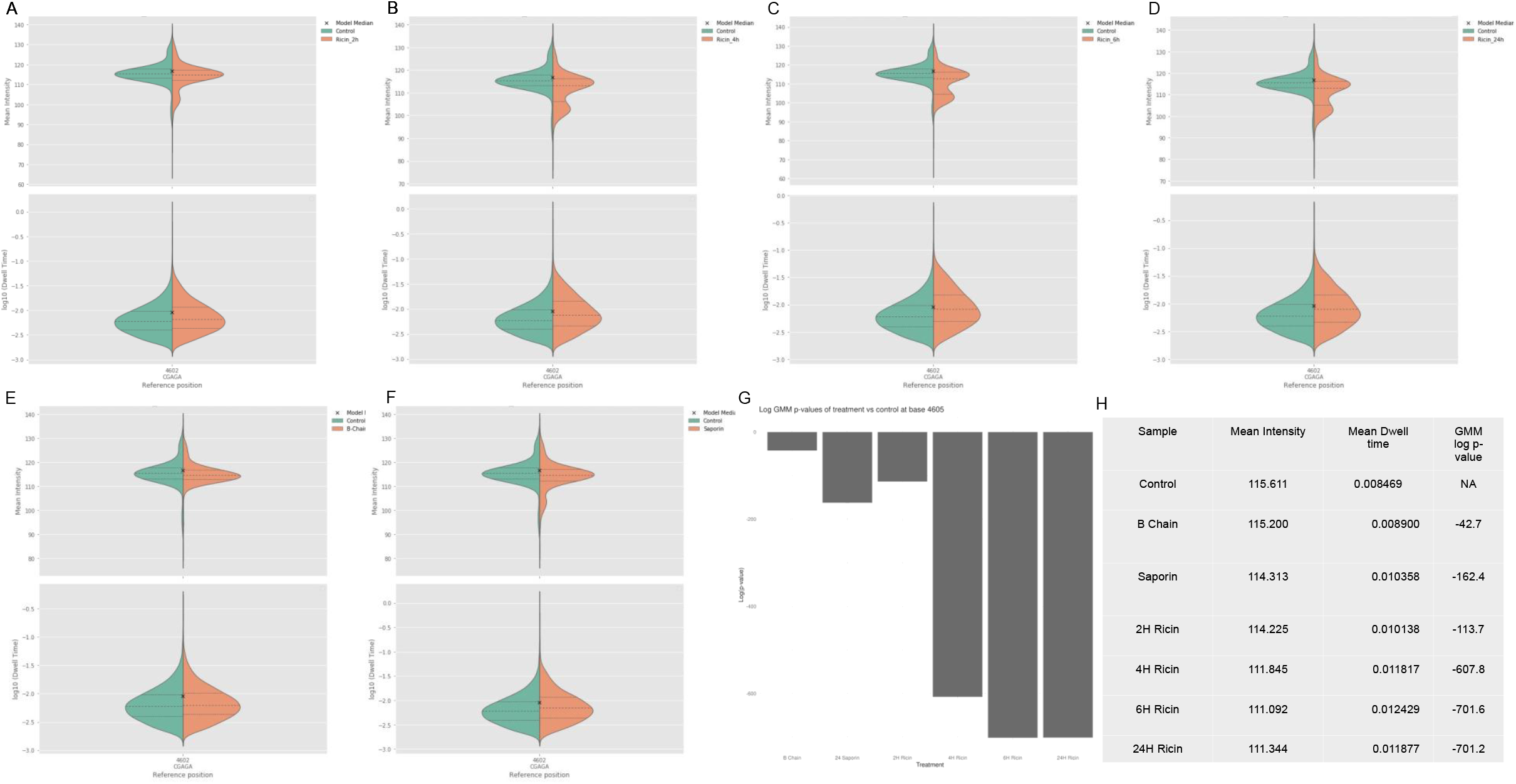
Charge density shown as a violin plot of the charge intensity per read at position 4605 on the 28s RNA determined by Nanocompore analysis of the direct RNA sequencing data (A-F). Shown are the dwell time and charge intensity at position 4605 from either A549 cells exposed to ricin for 2, 4, 6 and 24 hours (A to D), and RTB (E) and saporin (F). The control exposure is shown in green and the relevant experimental exposure in orange. The x indicates the mean, the middle-dashed line the medium and the outlying dotted lines the interquartile ranges. (G). To identify significant differences between the exposures, a pairwise comparison of the logged p-values of Gaussian mixture models was used. (H). Summary of the mean intensity, dwell time and the log p-value of the difference conditions.

The affect occurred not only at site 4605 but altered both charge and dwell time of nucleotides upstream and downstream, but no further than three nucleotides (Fig. 5). Assessment of how many nucleotides up and downstream of position A4605 were affected by depurination was performed by extending the window used by Nanocompore to measure the charge density and dwell time (Fig. 5). B-chain produced little effect compared to controls as was expected (Fig. 5A). A six-hour exposure to ricin resulted in the signal changes from the abasic site extending primarily across nucleotides 4603-4606 (Fig. 5B). Nucleotide position 4606, as with position 4605, showed a decrease in charge intensity, whereas position 4603 showed an increase in charge intensity. The was little change in charge density at position 4604, but an increase in dwell time, indicating a slower translocation speed, corroborating the effect of depurination observed previously^16^. See Supplementary Fig. 2 for violin plots of charge density and dwell time across all conditions.

**Fig. 5.**
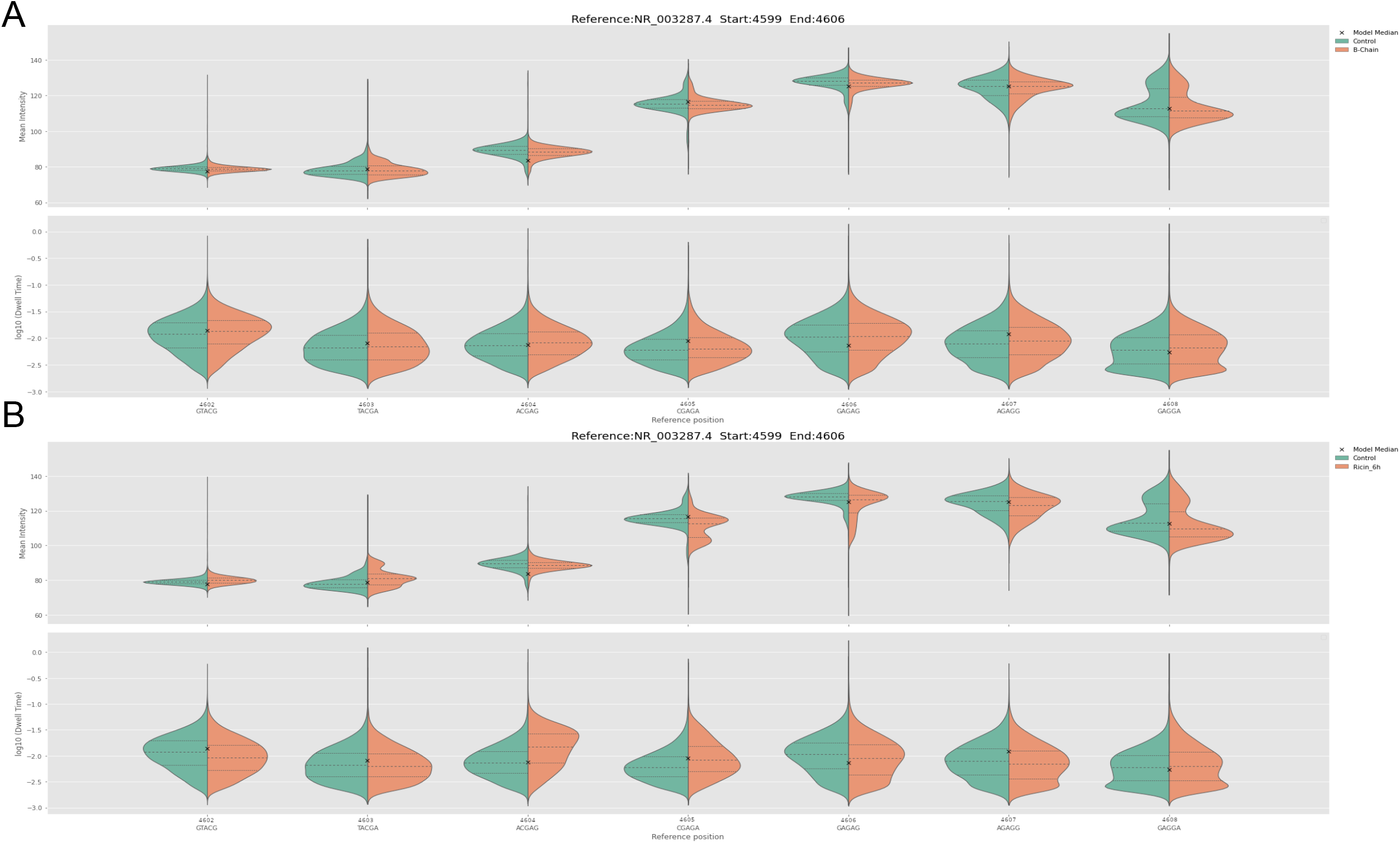
Nanocompore generated violin plots of charge density and dwell time of; (A). Control vs B-chain and (B). Control vs 6 hours ricin exposure. The control exposure is shown in green and the relevant experimental exposure in orange. The x indicates the mean, the middle-dashed line the medium and the outlying dotted lines the interquartile ranges. B-chain and controls gave similar profiles. For (A) and (B), the order of the violin plots represented the sequence around the ricin lop, starting with site 4602 (left) and extends to site 46018 (right). Ricin exposure caused a decrease in charge at sites 4605 and 4606, but an increase in charge at position 4603. Site 4605 and to a greater degree 4604, increased in dwell time in RNA from cells exposed to ricin.

### The decrease in charge intensity due to depurination at position A4605 mimics uracil in the ricin loop

To elucidate the mechanism behind the base calling of uracil at the apurunic 4605 in the ricin exposed cells and the associated lower charge intensity, charge intensities of the ricin loop were simulated using Nanocompore such that an A4605 was compared to an U4605 (Figs. 6A and 6B, respectively). A sample from ricin exposed cells would contain sequence reads that are either called as A4605 or U4605. These were matched, using Tombo^17^, to the actual recorded charge intensities in the ricin loop from control (predominately A4605) or from cells exposed to ricin (Figs. 6C and 6D, respectively). Tombo was used to superimpose the recorded charge intensities onto the predicted charge intensities from the reference sequence. The sequence reads from cells exposed to ricin were segregated to only those called as U4605, the reads mapping to A4605 were disregarded (Fig. 6D). The reference sequence was modified to U4605 and aligned to the actual charge intensity when the reads containing U4605 were considered in isolation.

**Fig. 6.**
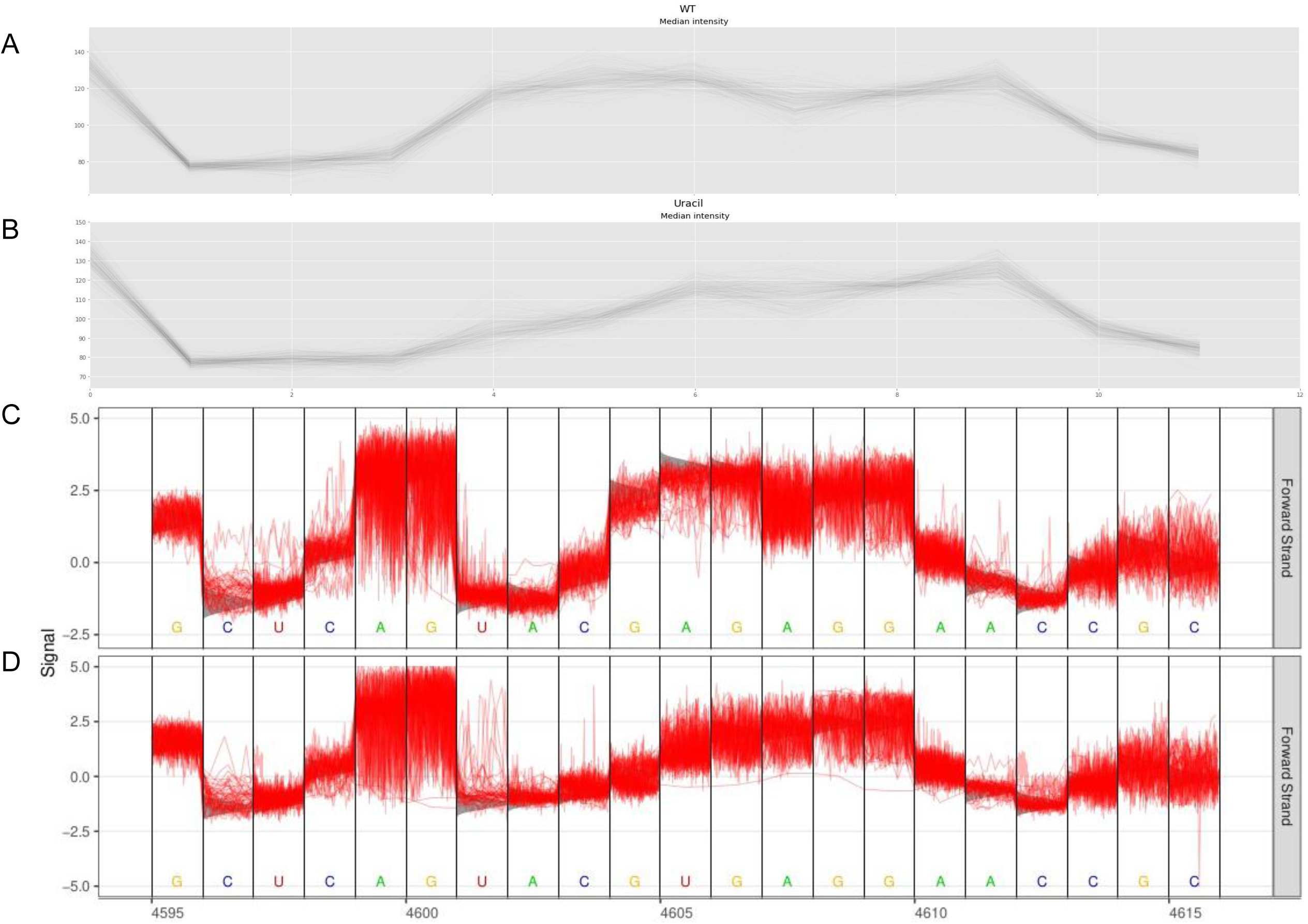
Comparison of Nanocompore median simulated charge intensity with the canonical ricin loop reference sequence (A), and sequence with the apurinic site replaced with an uracil (B). For A and B, the x-axis is the reference position, and the y-axis is charge intensity (C). Tombo output displaying the charge intensity of the ricin loop from control (non-exposed) A549 cells. (D). Tombo output for the ricin loop from cells exposed to ricin for 6 hours, mapped to a reference where the A4605 is altered to U4605. For C and D, the x-axis is the position of the nucleotide on the 28s RNA and y-axis is a normalised charge intensity of the actual (red) and predicted (grey) sequenced reads.

The simulated reads showed a marked decrease in charge intensity at base 4605 comparing A4605 (Fig. 6A) to U4605 (Fig. 6B). This paralleled the findings from the Nanocompore analysis (Fig. 5B) which showed a decrease in charge associated with an apurunic site at position 4605. See Supplementary Fig. 3 for simulations with cytosine and guanidine at position 4605. This suggested that Guppy was miscalling the apurunic site as an uracil. When viewing the charge intensities through Tombo (Fig. 6C), the control reads aligned closely with the predicted charge for A4605. Whereas, the immediate nucleotide upstream of site 4605 decreased in charge intensity with depurination of site 4605. This was observed for both the actual and predicted charge intensities. This identified the likely mechanism for Guppy base calling the apurunic site 4605 and supported the use of quantifying uracil calls as a proxy for depurination.

## Discussion

Utilising hyper-specific removal of adenine 4605 by ricin to only a single abasic site, we demonstrate the first detection of depurination with direct RNA sequencing on an ONT flow cell (using either the MinION or GridION sequencer). This modification was identified through base calling of direct RNA sequence data producing a transposition of A4605 to U4605 in depurinated reads (Fig. 3). To investigate why the modified base was identified as uracil and how this could be exploited as a way of detecting depurination and RIP activity, signal level comparison of RIP exposed, and unexposed reads was performed. The data indicated that the charge intensity and the dwell time were different at base 4605 from RNA obtained from cells exposed to ricin and saporin, compared to unexposed controls (Fig. 4, Fig. 5). This provided a likely explanation for Guppy base calling depurinated 4605 as uracil. To validate this, Nanocompore was used to simulate the 28s rRNA with U4605 and produced reads with a lower charge at the 4605 site versus A4605 (Fig. 6 A and B). Tombo produced a canonical model with expected charge intensities for a given sequence, which was compared against the recorded charge intensities. The analysis using Tombo, comparing U4605 and depurinated reads with A4605 and non depurinated reads, concurred with the Nanocompore simulated reads (Fig. 6 C and D). The lower charge associated with depurination at 4605 was the cause of miss-base calling as U4605, and gave credence to quantification of RIP induced depurination by measuring levels of U4605 with RIPpore.

While this investigation focused only on a single site of depurination, it provides the possibility of creating a basecaller model that is depurination aware. Software such as CURLCAKE (https://cb.csail.mit.edu/cb/curlcake/) can be used to generate a set of nucleotides covering each possible kmer^18^. These sequences can be synthesised, and randomly depurinated by decreasing the pH^16, 19^ of their storage solution. Sequencing of depurinated and unexposed samples would provide a dataset that could enable training of a depurination aware basecaller.

RIPpore allowed for quantification of RIP induced depurination and demonstrated the time dependent nature of the effect of ricin exposure on 28s ribosomes. This matched the observation using digital droplet PCR^15^. Our study found that after 24 hours of exposure, the proportion of depurinated ribosomes had begun to decrease. Whether this effect was due to cell death removing depurinated ribosomes or transcription of additional 28s rRNA to combat the loss of functional ribosomes is currently unclear. There is also the possibility that the cell was attempting to recover by creating additional ribosomal RNA, but was unable to translate the required proteins for correct ribosome folding which may then circumvent the action of ricin, resulting in increased non-depurinated reads^20^.

For creating and testing countermeasures to RIPs, such as ricin, direct RNA sequencing will be highly useful in quantifying the relative protection against depurination/ricin exposure a treatment provides. RIPpore provides a quantitative assessment of depurination from direct sequence read data. This technique would also be of benefit to those developing RIP based immunotoxins to assess off target ribosome inactivation across various cell types and targets. This method also creates an additional wealth of data on the epi-state of the 28s rRNA due to other modifications, which as of yet is a relatively unexplored field using direct RNA sequencing. This will enhance investigation into ribosomal regulation due to differing modifications in a wide variety of contexts.

## Methods

### Ricin Extraction

The ricin preparations used for this investigation were conducted at Dstl. Ricin was extracted from castor beans of *Ricin communis subsp. zanzibariensis* as per methodology previously published^21^.

### Assessment of ricin preparations by SDS electrophoresis

The purity of the ricin preparations was assessed by SDS PAGE using coomassie blue staining (GE Healthcare). Samples (1μL at 5 or 10 mg/mL) were mixed with a 1:1 volume of sample buffer Tris-HCl (125 mM) pH to 6.8, SDS (10%, w/v), glycerol (50%, v/v). Samples were heated at 95°C for 5 min prior to loading 1 μL onto the gel (polyacrylamide gradient (4-14%); GE Healthcare). Proteins were separated for 40 min before Coomassie blue staining for visualization, using manufacturer instructions (Calibrated Densitometer GS-800; Bio-Rad, Hemel Hempstead, UK). Densitometric scans were analyzed using Quantity One software (Bio-Rad). The value of the ricin band was determined as a percentage of the total protein in each lane, using the average density analysis tool.

### Cell culture

hACE-A549 cells were kindly provided by Olivier Schwartz^22^. Cells were cultured in 10% foetal bovine serum and Dulbecco’s Modified Eagle Medium at 37°C and 5% CO_2_. Cells were cultured to 80-90% confluency in a T75 before exposure.

### In Vitro Cytotoxicity Assay

A549 (86012804) cells were obtained from the European Collection of Animal Cell Cultures (ECACC) (Public Health England, Salisbury, UK). Cells were maintained in culture medium consisting of DMEM (Sigma Aldrich, Poole, UK) with 10% (v/v) foetal calf serum (Sigma Aldrich, Poole, UK), 1% penicillin, streptomycin solution containing 100 units/mL penicillin and 0.01 mg/mL streptomycin), and 1% (w/v) l-glutamine (Sigma Aldrich, Poole, UK) 2 mM. Cells were grown in T75 flasks in a humidified atmosphere of 5% CO_2_ in air at 37°C and removed from the flask surface using incubation with trypsin (0.05% w/v) (Sigma Aldrich, Poole, UK) containing EDTA (0.03% w/v) (Sigma Aldrich, Poole, UK) on achieving 70% confluency. The monolayer was allowed to adhere to the culture plates for 24h before addition of Ricin. For toxicity assessment, ricin toxin was diluted to 100 ng mL^-1^ in culture medium and filtered using a 0.2 m sterile filter before further dilution in culture medium and addition to the assay plate in triplicate. The plates were then incubated for 48 hours prior to the addition of 10 μL of Roche Cell Proliferation Reagent WST-1 (Sigma Aldrich, Poole,UK). After 3 hours the absorbance was read on a Thermo Multiskan plate reader (Thermo Fisher Scientific, Loughborough, UK) at 450nm to assess cell viability.

### RIP Exposure

All RIPs were handled with appropriate safety measures. Ricin was added to cell culture media of a volume appropriate to the cell culture plate or flask being used to create a solution of 1nM. Saporin was added as described by Rust et al, at 100 nM^23^ for 24 hours. B-chain was added at 1 nM for 6 hours. Ricin exposure lasted 2, 4, 6 and 24 hours. Cell culture media was then disposed of in sodium hypochlorite solution for inactivation. Care was taken not to generate aerosols at this step. Two PBS washes of the monolayer were performed to remove any remaining RIPs and culture media present. Waste PBS is disposed of in sodium hypochlorite solution.

### RIP Disposal

RIPs were deactivated by disposal into a solution of sodium hypochlorite with a minimum of 10,000 ppm active chlorine for 24 hours. Sodium hypochlorite and inactivated RIPs were disposed in accordance with local regulations.

### RNA Extraction

RNA was extracted using Trizol solution (Thermo Fisher) and optionally in conjunction with Phasemaker tubes (Thermo Fisher) according to manufacturer instructions.

### Quantifying RNA content

The RNA content of samples was measured using the Qubit RNA system (Thermo, Qubit RNA HS Assay Kit, Q32855). The Qubit working solution and standards were prepared as per manufacturer instructions. 2 μL of each sample was then added to a tube containing 198 μL of working solution and vortexed for 2-3 sec. All tubes were incubated at room temperature for 2 min. Samples were read using a Qubit 3.0 Fluorometer using the high sensitivity method.

### Targeted direct RNA oligos A and B with and without barcodes

The following oligos were synthesised from Eurofins with HPLC purification, and were used to replace the standard Oligos A and B in the direct RNA sequencing protocol. The last 10 nucleotides of Oligo B are the reverse complement of the last 10 nucleotides 3’ of 28s rRNA AGGGTTTGTC resulting in GACAAACCCT from NCBI accession: NR_003287.4. Unbarcoded reads may modify ONT’s oligo A and B as described in Oxford Nanopore Sequence specific direct RNA protocol DSS_9081_v2_revK_14Aug2019. Deeplexicon and Poreplex ^24, 25^ oligos were modified to replace the poly T tail with 28r rRNA sequence specific sequence.

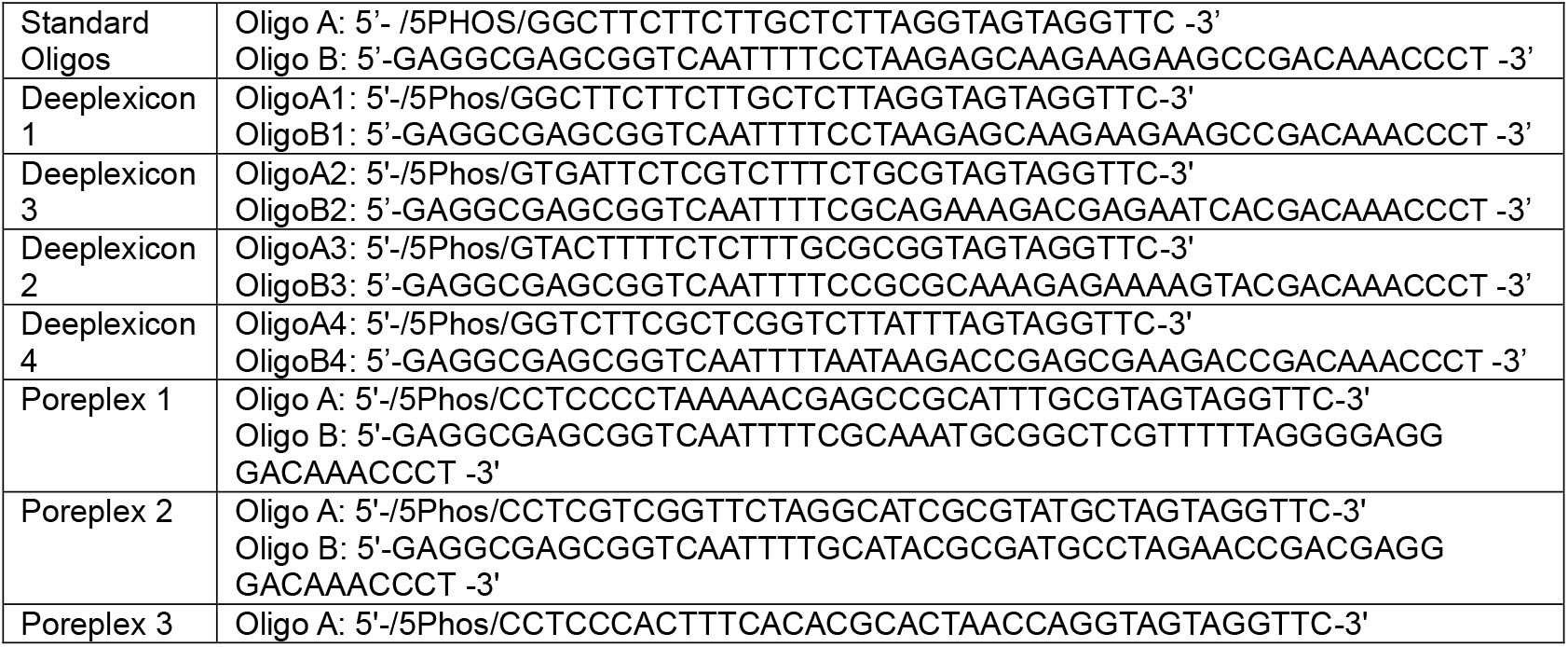
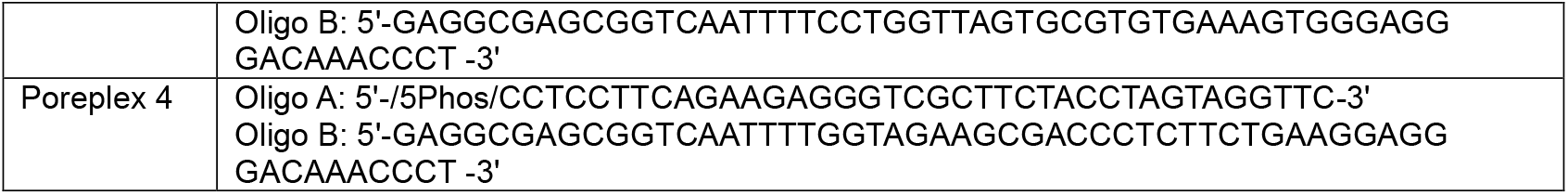

### Oxford Nanopore Library Preparation

Oxford Nanopore Sequence specific direct RNA protocol DSS_9081_v2_revK_14Aug2019 was used in this study. Note the reverse transcription step to linearise the 28s rRNA was performed as secondary structure inhibits translocation through the nanopore. For both barcoded and unbarcoded library preparations, 1.4 μM of oligos A and B were annealed 1:1 in 10 mM Tris-HCl pH 7.5, 50 mM NaCl as described in the direct RNA protocol. For each sample, whether barcoded or not, 2 μg of extracted RNA, split into two, 1 μg samples. Each sample proceeds through the Oxford Nanopore protocol as directed, until step 18, and eluted in 10 μl nuclease free water. All samples were pooled at step 20 and adjusted with the NEBNext Quick Ligation Reaction Buffer as needed for the volume of reverse transcribed RNA. The protocol then proceeds as normal.

### Oxford Nanopore Sequencing

Sequencing was undertaken on MinION MD106 flow cells, with SQK-RNA002 kit, and those kits appropriately selected in Minknow. Guppy version 4.4.1 was used for base calling. Older runs were re-basecalled with version 4.4.1 for parity.

### Demultiplexing

Poreplex^24^ was used to demultiplex individual samples using the options: –trim-adapter – barcoding –fast5. For one set of samples. DeePlexiCon^25^ was used with default options. Note any future direct RNA barcoding and demultiplexing tools which work in a similar method to the above tools will likely produce a satisfactory output.

### Quantification of depurination

Quantification of depurination was achieved using the following workflow:

1. Concatenate all fastqs in fastq_pass per sample
2. Map concatenated fastqs to NR_003287.4 using: minimap2 -ax splice -uf -k14 ref.fa reads.fq > aln.sam minimap2^26^ version 2.17-r941 was used
3. Sort and index sam file using Samtools Samtools^27^ version 1.9 was used
4. Run rippore.py to calculate per base nucleotide counts. By default, uses base 4605, if using a different 28s rRNA reference to NCBI: NR_003287.4, provide the correct base with -b option.

rippore.py -s sample.sam

Summary statistics and Figure 2 were generated in R version 4.0.2 and rippore.py with the -o option to create a CSV of nucleotide counts at a position.

### Raw Signal Analysis

Nanocompore^28^ version 1.0.3 was used to identify changes in charge intensity and dwell time at a range of locations on the 28s ribosome, primarily centered around site 4605. This analysis used the same NCBI NR_003287.4 as a reference. Nanocompore was also used to generate simulated reads of the 28s ribosome, both wildtype and modified to T4605. The Nanopolish SimReads function was used in its default settings, and no modification options enabled.

The references used were:

>WT

AGTACGAGAGGAACCG

>Uracil

AGTACGTGAGGAACCG

>Guanidine

AGTACGGGAGGAACCG

>Cytosine

AGTACGCGAGGAACCG

Tombo^17^ version 1.5.1 was used to generate aggregate plots of 50 random reads around the ricin loop, for WT and A4605, depurinated and T4605 to assess changes in read signal against Nanocompore simulated signals.

## Acknowledgements

We acknowledge the MRC DiMeN DTP for funding YR and AH as part of a PhD CASE studentship with dstl.

## Author contributions

YR and AH performed the ricin exposure and sequencing. AH and CH helped with cell culture and sequencing. HT assisted in testing and optimising direct RNA barcoding. The data was analysed by YR and JAH. JD, GC and JAH oversaw the project and provided supervision. RIPpore was written by YR. YR, AH and JAH wrote the manuscript with all other authors providing editorial corrections and approval of the final form.

## Data availability

Fast5 files available from SRA. BioProject PRJNA752594 (https://www.ncbi.nlm.nih.gov/bioproject/PRJNA752594).

RIPpore can be found at https://gitlab.com/yryan/rippore

## Supplementary Data

**Supplementary Table 1.** Overview of Flow cell and direct RNA reads. Flow cells show a range of 15,000 to 600,000 reads, with no discernible effect from barcoding on final number of reads.

| Flow cell | Total Reads | Samples Loaded |
| --- | --- | --- |
| Control 1 | 441184 | 1 |
| Control 2 | 79569 | 1 |
| Control 3 | 14500 | 1 |
| B-chain | 296000 | 4 |
| Ricin Cell 1 | 91437 | 3 |
| Ricin Cell 2 | 556362 | 4 |
| Ricin Cell 3 | 651134 | 4 |
| Poreplex Cell | 192358 | 4 |
| Saporin Cell | 308952 | 4 |

**Supplementary Table 2.**
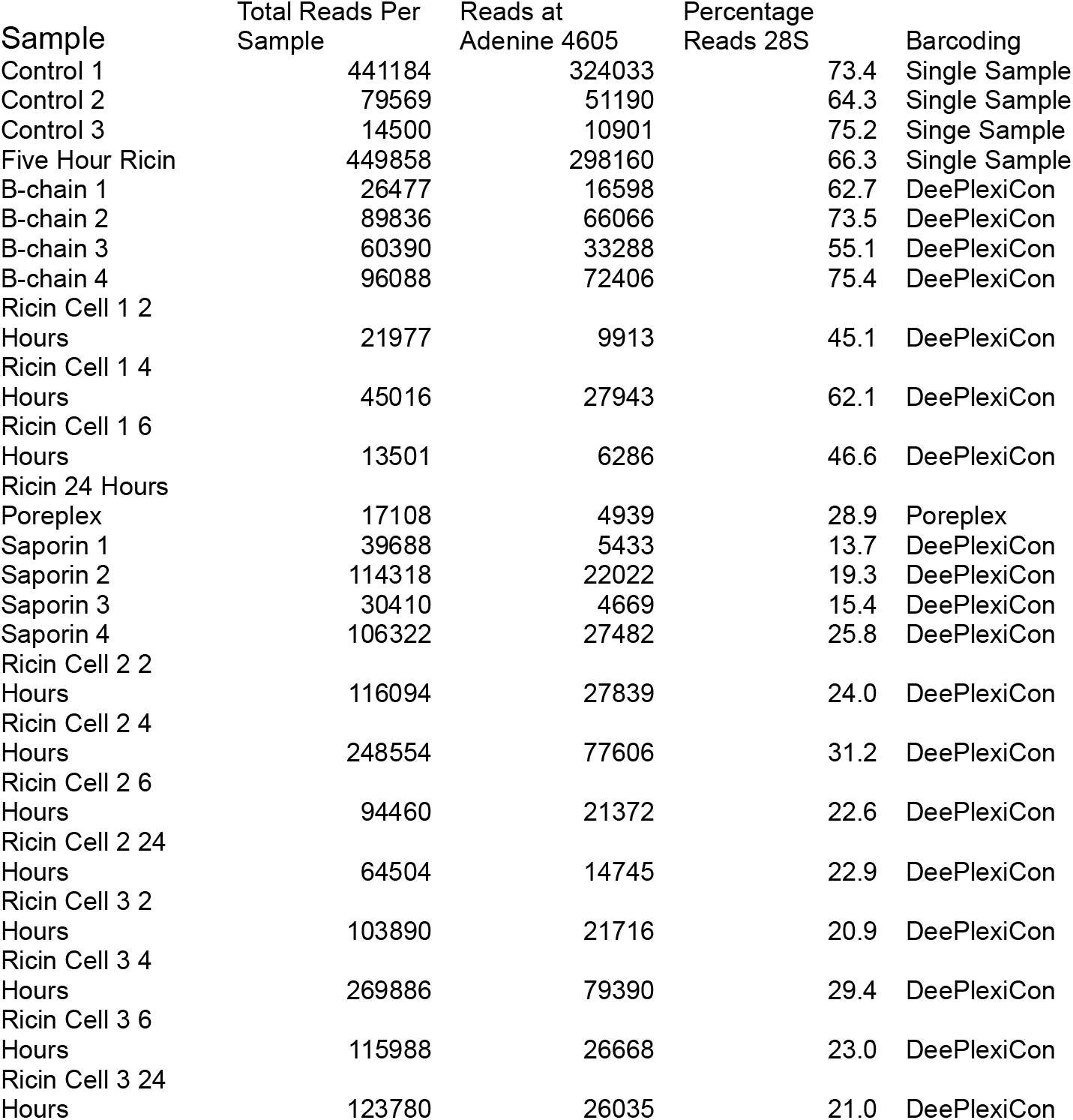
Overview of Sample Reads per sample and reads at RIP depurination site. Samples show generally the same order of magnitude of reads with two saporin samples, and one ricin sample in flow cells 2 and 3 being an order of magnitude lower than the remaining reads. The efficiency of the custom barcode ligation to 28S ribosomal reads has a high variance between 15 and 75% but an average of 41% as seen by percentage Reads 28S.

**Supplementary Fig. 1.**
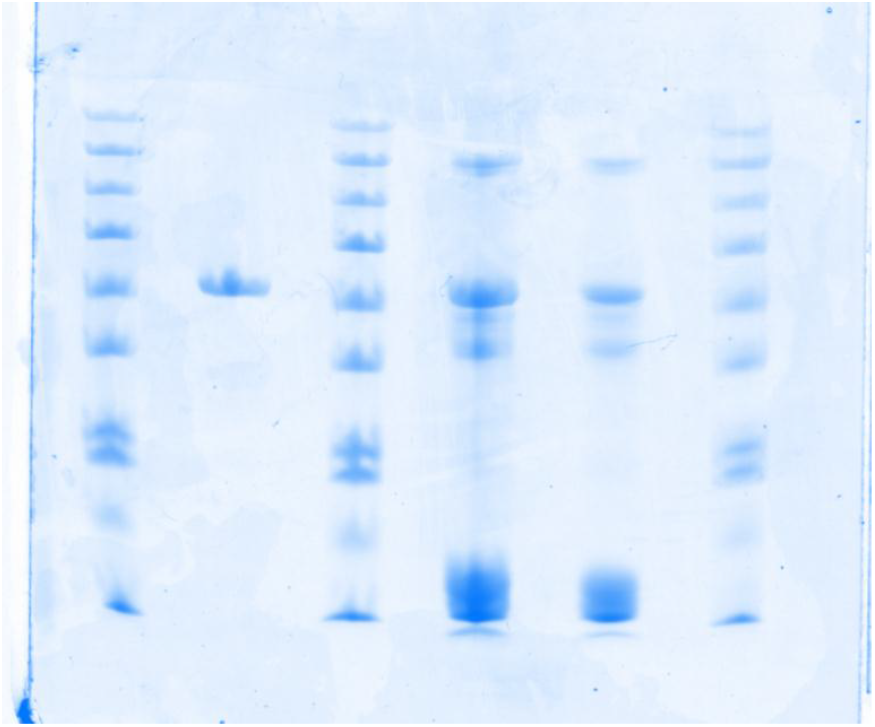
Unedited image of Fig. 2(A) in original orientation. Shows Coomassie blue analysis of crude (right) and purified (left) ricin toxin preparations alongside protein ladders (10-180 kDa).

**Supplementary Fig. 2.**
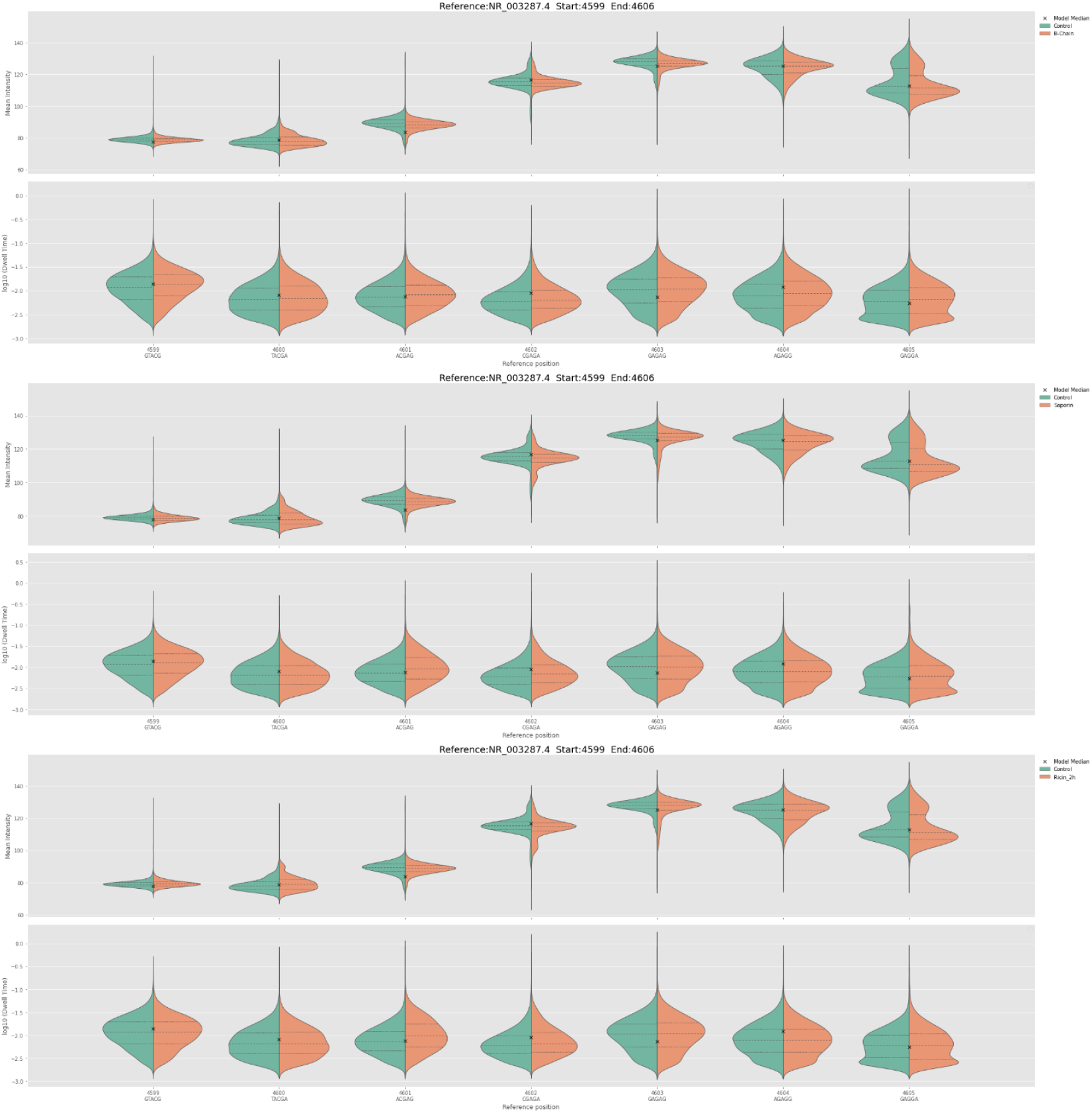
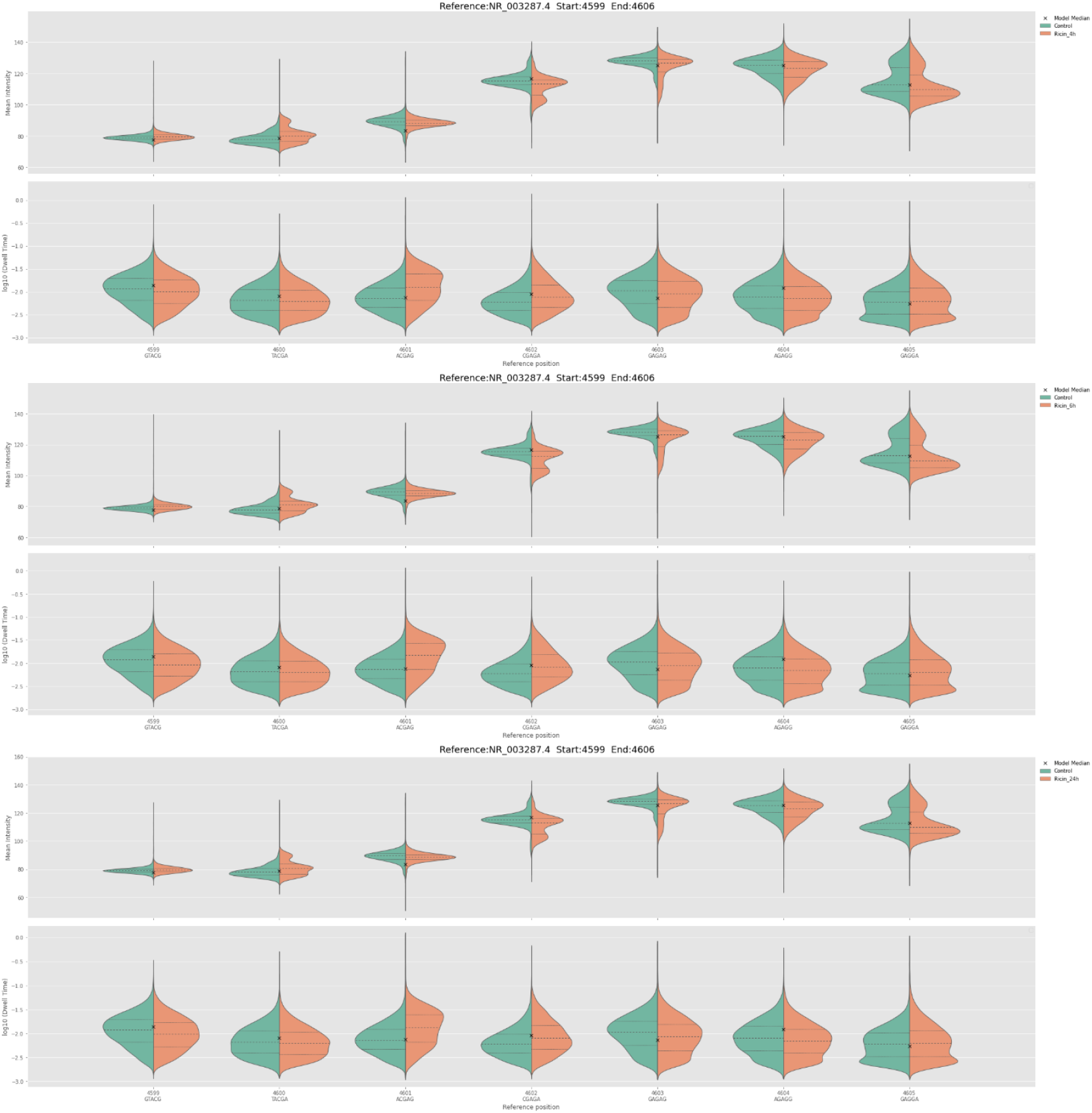
Fig. 5 Extended to All Conditions. shows time dependence of the effects on charge density and dwell time, with saporin and earlier timepoints showing more similarity to control reads, and later ricin exposed time points showing greater changes.

**Supplementary Fig. 3:**
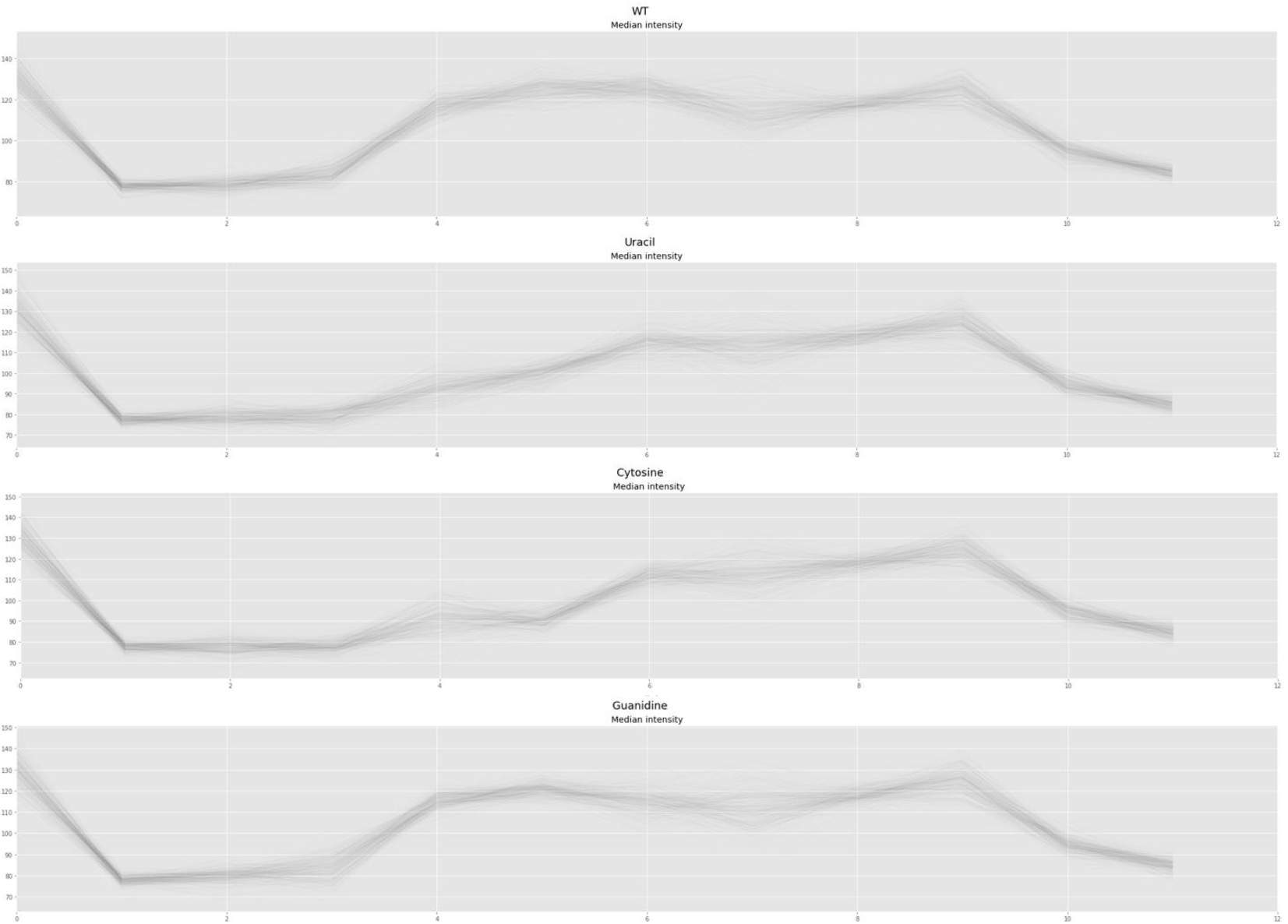
Fig. 6 extended with all four nucleotides. shows Nanocompore simulated charge intensities for reads with all four bases at base 4605. Cytosine has a decrease in charge similar to uracil but plateaus between positions 4 and 5, whereas uracil continues decreasing. Guanidine appears similar to adenine, but at position 5 peaks at around 120 charge intensity, whereas adenine peaks nearer 130.

